# Interdependence of CTL and NK cell cytotoxicity against melanoma cells

**DOI:** 10.1101/2020.06.14.150672

**Authors:** Kim S. Friedmann, Arne Knörck, Sabrina Cappello, Cora Hoxha, Gertrud Schwär, Sandra Iden, Ivan Bogeski, Carsten Kummerow, Eva C. Schwarz, Markus Hoth

## Abstract

CTL and NK cells recognize and eliminate cancer cells. However, immune evasion, down regulation of immune function by the tumor microenvironment, or resistance of cancer cells are a major problem. While CTL and NK cells are both important to eliminate cancer, most studies address them individually. In a new experimental human model, we analysed combined primary human CTL and NK cell cytotoxicity against the melanoma cell line SK-Mel-5. At high effector-to-target ratios, MART-1-specific CTL or NK cells eliminated SK-Mel-5 cells within 24 hours indicating that SK-Mel-5 cells are initially not resistant. However, at lower effector-to-target ratios, which resemble conditions of the immune contexture in human cancer, a significant number of SK-Mel-5 cells survived. Whereas CTL pre-exposure induced resistance in surviving SK-Mel-5 cells to subsequent CTL or NK cell cytotoxicity, NK cell pre-exposure induced resistance in surviving SK-Mel-5 cells to NK cells but not to MART-1 specific CTL. In contrast, there was even a slight enhancement of CTL cytotoxicity against SK-Mel-5 cells following NK cell pre-exposure. In all other combinations, resistance to subsequent cytotoxicity was higher, if melanoma cells were pre-exposed to larger numbers of CTL or NK cells. Increases in human leukocyte antigen class I expression correlated with resistance to NK cells, while reduction in MART-1 antigen expression correlated with reduced CTL cytotoxicity. CTL cytotoxicity was rescued beyond control levels by exogenous MART-1 antigen. This study quantifies the interdependence of CTL and NK cell cytotoxicity and may guide strategies for efficient CTL-NK cell anti-melanoma therapies.

**Key points summary:**

- Cytotoxic T lymphocytes (CTL) and natural killer (NK) cells eliminate cancer cells. CTL and NK work in parallel, but most studies address them individually.
- In a new human experimental model, antigen-specific CTL and NK cell cytotoxicity interdependence against melanoma is shown.
- Whereas high numbers of antigen-specific CTL and NK cells eliminate all melanoma cells, lower, more physiological numbers induce resistance, in case secondary CTL or NK cell exposure follow initial CTL cell exposure or if secondary NK cell exposure follows initial NK cell exposure; only if secondary CTL exposure follows initial NK cell exposure no resistance of melanoma but even a slight enhancement of cytotoxicity was observed.
- Alterations in HLA-I expression correlated with resistance to NK cells, while reduction in antigen expression correlated with reduced CTL cytotoxicity. CTL cytotoxicity was rescued beyond control levels by exogenous antigen.
- The results should help to better understand and optimize immune therapies against cancer.

**Graphical abstract:** 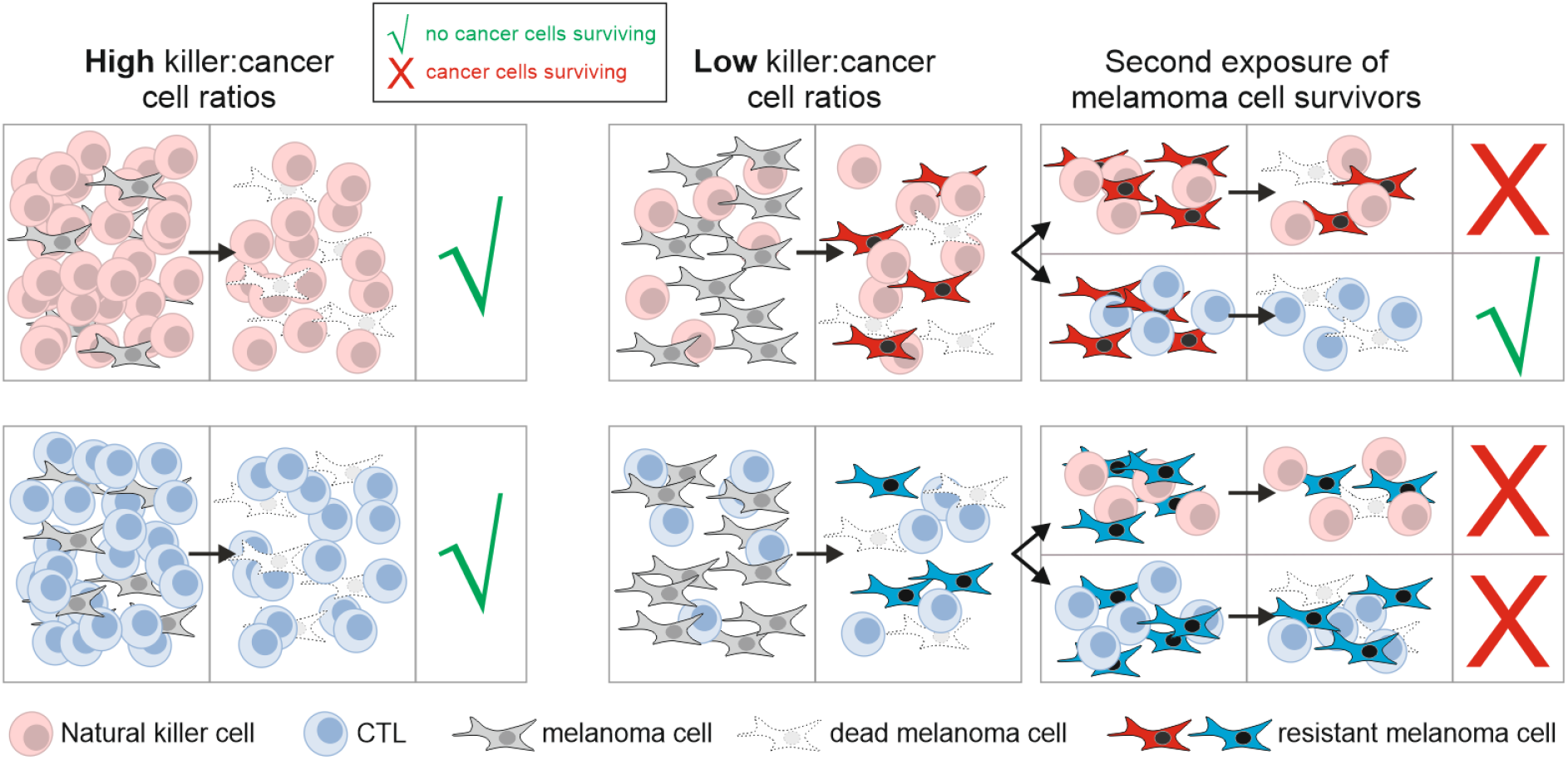

## Introduction

Cytotoxic T lymphocytes (CTL) and natural killer (NK) cells eliminate cancer cells in the human body. There is good evidence for a key role of CTL and NK cells in cancer immune surveillance. Already 20 years ago, Imai et al. found a clear correlation between natural lymphocyte cytotoxicity and cancer incidence in an 11-year follow up study in the general Japanese population (Imai *et al.*, 2000). In addition, Shankaran et al. reported strong arguments in favour of this hypothesis when they showed that lymphocytes protect against the development of carcinogen-induced sarcomas (Shankaran *et al.*, 2001). The authors, however, also showed that lymphocytes may select for cancer cells with decreased immunogenicity, which “explains the apparent paradox of tumor formation in immunologically intact individuals”. Another key finding by Galon et al. revealed that quantification of the type, density, and location of immune cells within colorectal cancer samples was a better predictor of patient survival than commonly used histopathological methods (Galon *et al.*, 2006). These and many other reports increased the interest in, as now called, the immune contexture or immunoscore of cancer.

The immune contexture of cancer is defined by Fridman et al. as the density, composition (including maturation), functional state (functionality) and organization (including location) of the leukocyte infiltrate in a tumor (Fridman *et al.*, 2011; Fridman *et al.*, 2017). The immune contexture of cancer is of course a key factor shaping the tumor microenvironment, as it influences the concentration of many soluble factors including cytokines, reactive oxygen species or Ca^2+^ (Frisch *et al.*, 2019). There is also increasing evidence that the immune contexture is correlated with the genomic landscape of cancer, as for instance recently shown in lung adenomatous premalignancy (Krysan *et al.*, 2019) or breast cancer (Tekpli *et al.*, 2019). In the latter study, the immune contexture defined by the genomic landscape was also correlated with cancer prognosis. According to the summary of data from a large series of publications (summarized in (Fridman *et al.*, 2017)), CD8^+^ T cell density in the tumor infiltrate and subtype composition are good prognostic markers for many different cancer types. For NK cells, there is also evidence that cytotoxicity correlates with cancer incidence. Besides a link between NK cell activity and colorectal cancer (Jobin *et al.*, 2017) or prostate cancer (Kastelan *et al.*, 1997) incidence, Barry et al. showed that the NK cell frequency correlates with the abundance of protective dendritic cell in human cancers including melanoma and with overall survival (Barry *et al.*, 2018). A recent bioinformatics approach on RNA-seq data revealed an improved survival rate for patients with metastatic cutaneous melanoma in case tumours showed signs of NK cell infiltration (Cursons *et al.*, 2019). Together these examples stress the necessity to analyse the interplay of cancer with immune cells including CTL and NK cells.

To fight a tumor, CTL or NK cells form a close contact with cancer cells, called immunological synapse (IS). By direct contact and cytokine etc. release, CTL, NK and other immune cells interact with and influence tumours in many ways, often referred to as immunoediting of tumours. This includes the elimination of a tumor as the successful version of immunosurveillance, but it also includes cancer-immune equilibrium, and may also induce the escape of tumours from the immune system in both natural and therapeutical cancer strategies (Muenst *et al.*, 2016).

Immune evasion of cancer is a severe problem that limits CTL and NK cell immune responses in the human body against many cancers. Even worse, cancer hijacks certain immune functions for its survival or growth (Hanahan & Weinberg, 2011). To understand immune evasion or resistance of cancer is also of importance to optimize immune responses in the human body through drug-based therapy. In their recent review, Garner and de Visser state that: “[…] major challenges hinder the progress of immuno-oncology, including a lack of insight into the optimal treatment combinations to prevent or revert resistance to immunomodulatory strategies” (Garner & de Visser, 2020). Personalized immunotherapy should integrate CTL and NK cell concepts in the future (Rosenberg & Huang, 2018) stressing the role for studying combined CTL-NK cell cytotoxicity.

Malignant melanoma represents a skin cancer with high mortality rate and increasing incidence worldwide (Schadendorf *et al.*, 2018). High ultraviolet radiation (UVR) exposure due do chronic sun-bathing drives mutations in the Trp53 tumor suppressor, thereby accelerating BRAF-dependent melanoma induction (Viros *et al.*, 2014). Recently, UVR mutation signatures have been linked to patient survival (Trucco *et al.*, 2019). Due to its poor response to many of the standard tumor therapies including radiotherapy and chemotherapy, prior to 2010 treatment options were very limited. The past decade, however, has spawned an enormous evolution in melanoma therapy, bringing both targeted and immunotherapy approaches to clinical practice (Jenkins & Fisher, 2020). Patients with mutations in the MAPK pathway may benefit from new molecular targeted strategies, directed against oncogenic BRAF and/or MEK signalling. Moreover, melanoma is a highly immunogenic cancer, which has raised great interest in targeting the immune contexture of this cancer. At physiologic conditions, endogenous T cell-driven immune checkpoints control self-tolerance, thereby preventing autoimmunity. Blocking key checkpoint molecules, such as the co-inhibitory receptors CTL Antigen-4 (CTLA-4), the programmed cell death protein 1 (PD-1) or its ligand PD-L1, can activate CTL to attack cancer. This discovery has led to a breakthrough in developing new cancer treatments (Ribas & Wolchok, 2018). The efficacy of monoclonal antibodies targeting these checkpoint entities has first been demonstrated in melanoma (Hodi *et al.*, 2010; Brahmer *et al.*, 2012).

Meanwhile, immune checkpoint blockade has not only evolved as first-line treatment strategy for patients with advanced and metastatic melanoma (Jenkins & Fisher, 2020), but is also used as effective immunotherapy for other cancers (Ribas & Wolchok, 2018). Despite the great advances that immunotherapy brought to oncological practice, it is important to note that many patients do not respond, while others relapse, suggesting yet unknown innate or acquired resistance mechanisms (O’Donnell *et al.*, 2019). Paradoxically, despite eliciting strong immune responses, melanoma cells can frequently evade immune surveillance likely due to their high phenotypic plasticity, which is considered one cause for treatment failure and/or resistance. Melanoma cells undergoing “phenotypic switching”, e.g. in response to inflammatory mediators, often display considerable non-genomic heterogeneity, suggesting that cancer cell plasticity is, at least in part, a dynamic response to microenvironmental factors (Holzel & Tuting, 2016).

While the importance of CTL targeting in immunotherapy is in general well-established, there are also increasing interests and efforts to target NK cells for treating melanoma (Lorenzo-Herrero *et al.*, 2018; Cursons *et al.*, 2019). In cases where melanoma cells have escaped CTL-mediated elimination, NK cell-based immunotherapy may reflect an alternative treatment option (Tarazona *et al.*, 2015). However, the interdependencies of CTL and NK cells in cancer cell cytotoxicity are poorly understood, as are their potential roles in immunotherapy resistance mechanisms.

To gain mechanistic insight into CTL, NK and cancer cell interactions, our study introduces a simple assay to quantify the combined efficiency of human CTL and NK cells against melanoma (and potentially also against other cancer cell types). We pre-exposed melanoma cells to CTL or NK cells and analysed the susceptibility of surviving melanoma cells to fresh CTL or NK cells. Conditions that do not allow complete eradication of cancer cells resemble conditions of an inefficient immune response during cancer development. We found a remarkable resistance of CTL-pre-exposed melanoma cells towards subsequent CTL and NK cell-driven cytotoxicity, whereas NK-pre-exposed melanoma cells showed a differential resistance phenotype towards CTL and NK cell-driven cytotoxicity, with implications for development of future immunotherapies.

## Methods

### Ethical approval

Research was approved by the local ethic committee (84/15; Prof. Dr. Rettig-Stürmer). The local blood bank within the Institute of Clinical Hemostaseology and Transfusion Medicine at Saarland University Medical Center provided leukocyte reduction system (LRS) chambers, a by-product of platelet collection from healthy blood donors. All blood donors provided written consent to use their blood for research purposes.

### Cells

T2 cells and lymphoblastoid cell lines were kindly provided by Dr. Frank Neumann (José Carreras Center for Immuno and Gene Therapy, Saarland University, Homburg, Germany) and cultured in RPMI 1640 medium (Thermo Fisher Scientific) supplemented with 10% FBS and 1% penicillin–streptomycin (Thermo Fisher Scientific). SK-Mel-5 cells (ATCC^®^ HTB-70™) were purchased from ATCC, a second batch of cells was kindly provided by the Department of Dermatology, Venerology and Allergology (University Hospital of the Saarland, Homburg, Germany), originally purchased from CLS (330157). All experiments except one, for which CLS was the source as indicated in the figure legend, were performed with SK-Mel-5 from ATCC. Other melanoma cell lines are from the following sources: 1205Lu (ATCC^®^ CRL2806™, provided by M. Herlyn, WISTAR Institute, Philadelphia), 451Lu (ATCC^®^ CRL2813™, provided by M. Herlyn, WISTAR Institute, Philadelphia), SK-Mel-28 (ATCC^®^ HTB-72™, provided by T. Vogt, Dermatology, Homburg), MeWo (ATCC^®^ HTB-65™, provided by T. Vogt, Dermatology, Homburg) and MelJuso (DSMZ ACC74, provided by T. Vogt, Dermatology, Homburg). SK-Mel-5 and other melanoma cells lines cells were maintained in Eagle's Minimum Essential Medium supplemented with 10% FBS and 1% penicillin–streptomycin at 37°C and 5% CO2, split twice a week and kept in culture no longer than 3 months.

Human peripheral blood mononuclear cells (PBMCs) were isolated from healthy donors after routine platelet-apheresis using LRS chambers of Trima Accel devices (Institute of Clinical Haematology and Transfusion Medicine, Homburg). For PBMC isolation, density gradient centrifugation using Lymphocyte Separation Medium 1077 (PromoCell) was carried out as described before (Knorck *et al.*, 2018). Primary NK cells were isolated from PBMC using Dynabead® Untouched™ NK cell isolation kits (Thermo Fisher Scientific) according to the manufacturer’s instructions as described previously (Backes *et al.*, 2018). NK cells were cultured in AIM-V medium supplemented with 10% FBS and 50 ng/ml IL-2.

### Generation of MART-1-specific CD8^+^ T-cell clones

A MART-1 specific stimulation of naïve CD8^+^ T-cells of an HLA-A2^+^ donor was carried out as described previously (Wolfl & Greenberg, 2014). Briefly, immature DC were differentiated from monocytes isolated by plastic adherence. Monocytes were stimulated with IL-4 and GM-CSF for 72 h followed by the addition of IL-4, LPS, IFNγ and MART-1 peptide and incubation for 16 h to induce the generation of mature MART-1-presenting DC. Naïve CD8^+^ T-cells were isolated from autologous PBMC using “Naive CD8^+^ T Cell Isolation Kit” (Miltenyi Biotec) in parallel to the addition of MART-1 to the immature DC fraction. Mature DC were irradiated with 30 Gy and co-incubated with naïve CD8^+^ T-cells in Cellgro DC medium supplemented with 5% human serum. On the same day IL-21 was added, whereas IL-7 and IL-15 were applied at day 3, 5, and 7. After 10 days of co-incubation MART-1-specific stimulation of CD8^+^ T-cells was stopped. CD8^+^ T-cells were re-stimulated with MART-1-loaded autologous PBMC (irradiated with 30 Gy) for 6 h to induce IFN-γ secretion. Afterwards antigen specific CD8^+^ T cells were isolated using IFN-γ secretion assay (Miltenyi Biotec). Single cells were seeded into individual wells (1 cell/200 μl in each well) in RPMI 1640 supplemented with 10% human serum, 1% penicillin-streptomycin, 30 ng/mL anti-CD3 antibody (OKT3, Biolegend), 25 ng/mL IL-2, 5 × 10^4^ heterologous PBMC/well (mix of 2-3 donors, irradiated with 30 Gy) and 5 × 10^4^ cells/well of lymphoblastoid cell lines (mix of 2 donors, irradiated with 120 Gy) in 96-well U-bottom plates. After 7 days, 50 μl of RPMI1640 supplemented with 10% human serum, 1% penicillin-streptomycin and 125 ng/mL IL-2 were added to each well. After another week of incubation, proliferating CD8^+^ T cells were transferred into 25 cm^2^ cell culture flasks filled with 20 ml of RPMI 1640 supplemented with 10% FBS, 1% penicillin-streptomycin, 30 ng/mL anti-CD3 antibody (OKT3), 25 × 10^6^ PBMC (mix of 2-3 donors, irradiated at 30 Gy) and 5 × 10^6^ cells of lymphoblastoid cell lines (mix of 2 donors, irradiated at 120 Gy) for expansion of CD8^+^ T-cell clone populations. At day 1, 3, 5, 8, and 11, 30 ng/mL IL-2 and 2 ng/mL IL-15 were added. Finally, antigen specificity was assessed using MART-1-specific dextramers in flow cytometry and antigen specific cytotoxicity was analysed using real-time killing assays (Kummerow *et al.*, 2014). Antigen-specific clones were frozen in aliquots. Experiments were performed at day 11–14 after thawing and expansion of clonal populations. CTL-MART-1 clone 3 (CTL-M3) was already used in another study (Hart *et al.*, 2019).

### Reagents

Calcein-AM and CountBright Absolute Counting Beads and DiOC_18_ were purchased from Thermo Fisher Scientific. The following antibodies were used for flow cytometry and stimulation: FITC-labelled anti-HLA-A2 (BB7.2, Biolegend, 1:40), AlexaFluor® 647-labeled anti-HLA-A,B,C (W6/32, Biolegend, 1:40), FITC-labelled anti-CD8 (SK1, Biolegend, 1:50), Ultra-LEAF anti-CD3 (OKT3, Biolegend, 30 ng/mL), APC-labelled anti-MART-1 (ELAGIGILTV) dextramers (Immudex, 1:5), APC-labelled A*0201 dextramer negative control (Immudex, 1:5). Antibodies for western blot: anti-γ-tubulin (Sigma; 1:1000), anti-GAPDH (Cell Signaling; 1:2000), anti-MART-1 (Agilent/Dako, 1:1000, kindly provided by the Department of Dermatology, Venerology and Allergology (University Hospital of the Saarland, Homburg, Germany)). 7-AAD viability staining solution was from Biolegend. MART-1 peptide (MART-1_26-35A27L_) was purchased from JPT. Dulbecco’s PBS, 1% penicillin-streptomycin, IL-2, LPS, RPMI 1640, Eagle’s Minimum Essential, AIM-V and CTS-AIM-V medium were from Thermo Fisher Scientific. IL-7, IL-15, IFNγ and IL-4 were from Peprotech. Cellgro DC medium was from CellGenix. GM-CSF was purchased from Gentaur. All other reagents were from Sigma-Aldrich.

### Pre-exposure of SK-Mel-5 to CTL-M3/NK cells

SK-Mel-5 cells were cultured in minimum essential media (MEM) supplemented with 10% FBS in 75 cm^2^ cell culture flasks (1 × 10^6^/flask). Immediately before co-culture, NK and CTL-M3 were irradiated with 30 Gy to prevent proliferation and were added at different effector-to-target ratios (indicated in text) to SK-Mel-5 cells. After 3-4 days, supernatant was disposed and after a PBS washing step, remaining SK-Mel-5 cells were harvested for subsequent experiments.

### Western blot

Cell pellets were frozen at −80 °C. After thawing, cells were lysed in lysis buffer (150 mM NaCl, 1 % Triton X-100, 0.5 % NP-40, 10 mM Tris (pH 7.4) supplemented with protease inhibitors (complete, EDTA-free; Roche) and 0.1 μl Benzonase (Sigma-Aldrich). Protein concentration of lysates was quantified using “Pierce BCA (bicinchoninic acid) Protein Assay Kit”. Denaturation was carried out in Laemmli buffer at 90 °C for 5 min. 75 μg of total proteins were separated by 15 % SDS–polyacrylamide gel electrophoresis and afterwards transferred onto polyvinylidene difluoride membranes using a transblot transfer chamber (X-Cell SureLock™, Invitrogen Novex Mini-cell, or Mini-PROTEAN Tetra Cell, Biorad). Western blots were probed with anti-MART-1 antibodies (1:1000). For the detection of reference genes, blots were probed with anti-γ-tubulin antibodies (1:1000) or anti-GAPDH antibodies (1:2000). Signals were developed in BioRad imaging system by using ECL solution (Pierce, Thermo Scientific). Densiometric analysis was carried out using the software ImageLab 5.2.1. and Excel. Expression of protein was normalized to the reference proteins γ-tubulin or GAPDH.

### Flow cytometry

0.5 × 10^6^ cells were washed in FACS buffer (PBS supplemented with 0.5% BSA). Afterwards cells were stained in 100 μl FACS buffer supplemented with corresponding antibodies and kept in the dark at room temperature for 20 min. After two subsequent washing steps in FACS buffer, the pellet was re-suspended in 200 μl FACS buffer and analysed. For dextramer staining, 0.5 × 10^6^ CTL-M3 were washed once in dextramer buffer (PBS supplemented with 5 % FBS) and re-suspended in 50 μl dextramer buffer. Staining procedure was carried out by adding 10 μl of dextramers and incubation of 10 min in the dark at room temperature. Subsequently, anti-CD8 antibodies were added and cells were kept in the dark at 4°C for 20 min. After two washing steps cells were resuspended in 400 μl dextramer buffer and analysed in BD FACSVerse™ Flow Cytometer (BD Biosciences). Data analysis was performed using FlowJo (X 10.0.7).

### Long term (> 24 hours) killing experiments

For each sample 5 × 10^4^ SK-Mel-5 cells were seeded in a well of a 48-well plate and incubated in 320 μl MEM supplemented with 10 % FBS for 6 h to facilitate adhesion. DiOC_18_ (Live Dead Cell-Mediated Cytotoxicity Kit, L7010 (Thermo Fisher Scientific)) was diluted 1:50 in MEM supplemented with 10 % FBS, and 80 μl of DiOC_18_ staining solution was added to each well (final concentration of DiOC_18_ was 12 μM). After overnight incubation the staining solution was removed, cells were rinsed in AIM-V medium supplemented with 10% FBS and incubated in 320 μl AIM-V medium supplemented with 10 % FBS for 2 h. 5 × 10^3^ – 1 × 10^6^ NK or CTL-M3 effector cells were added at the indicated effector-to-target ratios of 0.1:1 - 20:1 in a total volume of 80 μl AIM-V medium supplemented with 10 % FBS. After 24 h of co-incubation pictures were taken with a high-content imaging system (ImageXpress, Molecular Devices). Subsequently, cells at the bottom of each well were resuspended and collected in a 5 ml (12 × 75 mm) tube, wells were washed once with PBS supplemented with 0.5% BSA to collect remaining cells. Cells were centrifuged 5 min at 300 × g and resuspended in 450 μl PBS supplemented with 0.5% BSA. 50 μl of CountBright Absolute Counting Beads (C36950, Thermo Fisher Scientific) and 5 μl 7-AAD (Biolegend) were added. After 10 min, analysis was performed on a FACS ARIA III Flow Cytometer (BD Biosciences). 2 × 10^3^ events of the CountBright Absolute Counting Beads population were recorded per sample. Each condition was prepared and recorded in triplicates.

### Short-term (4 hours) real-time killing assay

To quantify the cytotoxicity of CTL-M3 and primary NK cells, a real-time killing assay was carried out as described before (Kummerow *et al.*, 2014). Briefly, target cells (T2 or SK-Mel-5 cells) were stained with 500 nM Calcein-AM in AIM-V medium supplemented with 10 mM HEPES. In case of T2-killing or for rescue experiments with SK-Mel-5, cells were loaded initially in AIM-V medium (supplemented with 10 % FBS) with 0.5 μg MART-1 peptide in low attachments cell culture plates for 1.5 h. After calcein staining, 2.5 × 10^4^ target cells were pipetted per well into 96-well black plates with clear-bottom (VWR/Corning) and kept in the dark for 20 min at room temperature. After settling down of target cells, effector cells (CTL-M3 or NK cells) were added cautiously at the indicated effector to target ratio and killing was measured in a Genios Pro (Tecan) reader using bottom reading function at 37 °C. Maximal killing rates were calculated as the maximum increase between two subsequently measured time points. Maximum target cell lysis was quantified at 240 min.

### Apoptosis-necrosis assay with Casper-GR

The apoptosis-necrosis assay with Casper-GR was essentially carried out as described before (Backes *et al.*, 2018). SK-Mel-5 cells were transiently transfected using the jetOptimus transfection reagent according to the manufacturer’s instructions in MEM supplemented with 10 % FBS in a 25 cm^2^ cell culture flask. The pCasper3-GR vector (Evrogen) was used (2μg/per bottle). After 6 h the medium was changed to MEM supplemented with 10 % FBS + 1 % penicillin-streptomycin + 0.2 μg/ml puromycin. Cells were incubated for 30 h, washed in PBS + BSA, and pCasper^+^ (GFP^+^ RFP^+^) cells were sorted on a FACS ARIA III sorter (BD Biosciences). pCasper^+^ SK-Mel-5 were incubated in MEM supplemented with 10 % FBS + 1 % penicillin-streptomycin + 0.2 μg/ml puromycin overnight. 2 × 10^3^ pCasper^+^ SK-Mel-5 cells per sample were resuspended in 80 μl of CTS AIM-V Medium without phenol red supplemented with 10 % FBS and then seeded in a well of a 384-well black plate. After 2 h resting in the incubator, CTL-M3 or NK cells were added in 20 μl CTS AIM-V medium without phenol red supplemented with 10 % FBS at the indicated effector-to-target ratios and cytotoxicity was analysed with the high-content imaging system (ImageXpress, Molecular Devices). Semi-automated analysis was performed using Imaris (Bitplane), ImageJ and Excel.

### Statistics

Data are presented as the mean ± SD (n = number of experiments) if not stated otherwise. Gaussian distribution was tested using D’Agostino & Pearson normality test. If not stated otherwise, one-way/two-way ANOVA or Kruskal-Wallis test were used to test for significance: *P < 0.05, **P < 0.01, ***P < 0.001. Statistics were calculated using Prism 7 software (GraphPad Software, La Jolla, CA, USA).

## Results

### Cytotoxic efficiency of MART-1-specific human CTL clones against MART-1-loaded target cells and melanoma cells

The protein MART-1, or Melan-A, is frequently overexpressed in melanoma. The optimal length of the immunodominant peptide was located to the decapeptide MART-1_26-35_ (Romero *et al.*, 1997), which is recognized by HLA-A2-restricted lymphocytes (Kawakami *et al.*, 1994). A change in position 2 from alanine to leucin results in the mutant MART-1_26-35A27L_ which allows a more stable HLA-A2-antigen binding and an increased CTL immune response (Romero *et al.*, 1997). In addition, many T cells from the naïve repertoire express T cell receptors (TCR) specific for MART-1_26-35A27L_ (Zippelius *et al.*, 2002).

To analyse the combined cytotoxicity of CTL and NK cells against MART-1-positive melanoma target cells, we first generated MART-1-specific CTL clones from primary human PBMC. As the MART-1 antigen MART-1_26-35A27L_ is specific for the MHCI serotype HLA-A2, we chose an HLA-A2 positive blood donor to generate MART-1-specific CTL following a modified protocol by Wölfl and Greenberg (Wolfl & Greenberg, 2014). We screened and expanded 168 CTL clones from 5 independent cloning approaches. To test their cytotoxicity, we used T2 and SK-Mel-5 cells as target cells. T2 cells are hybrids of a human T- and B-cell line and are TAP (transporter associated with antigen processing)-deficient but HLA-A2 positive (Salter & Cresswell, 1986). They were chosen because, considering their TAP deficiency, they cannot transport intrinsic antigens to the cell surface and can thus easily be loaded with different concentrations of exogenous HLA-A2 specific antigens. SK-Mel-5 are human melanoma cells and were chosen because they express large amounts of MART-1 antigen (Du *et al.*, 2003) and are susceptible to both, CTL cytotoxicity (Sugita *et al.*, 1996) and NK cell cytotoxicity (Lee *et al.*, 2011). We confirmed high MART-1 expression of SK-Mel-5 cells by quantification against γ-tubulin expression in comparison to MelJuso, MeWo, SK-Mel-28, 451Lu and 1205Lu melanoma cells (Fig. 1A, B).

**Fig. 1:**
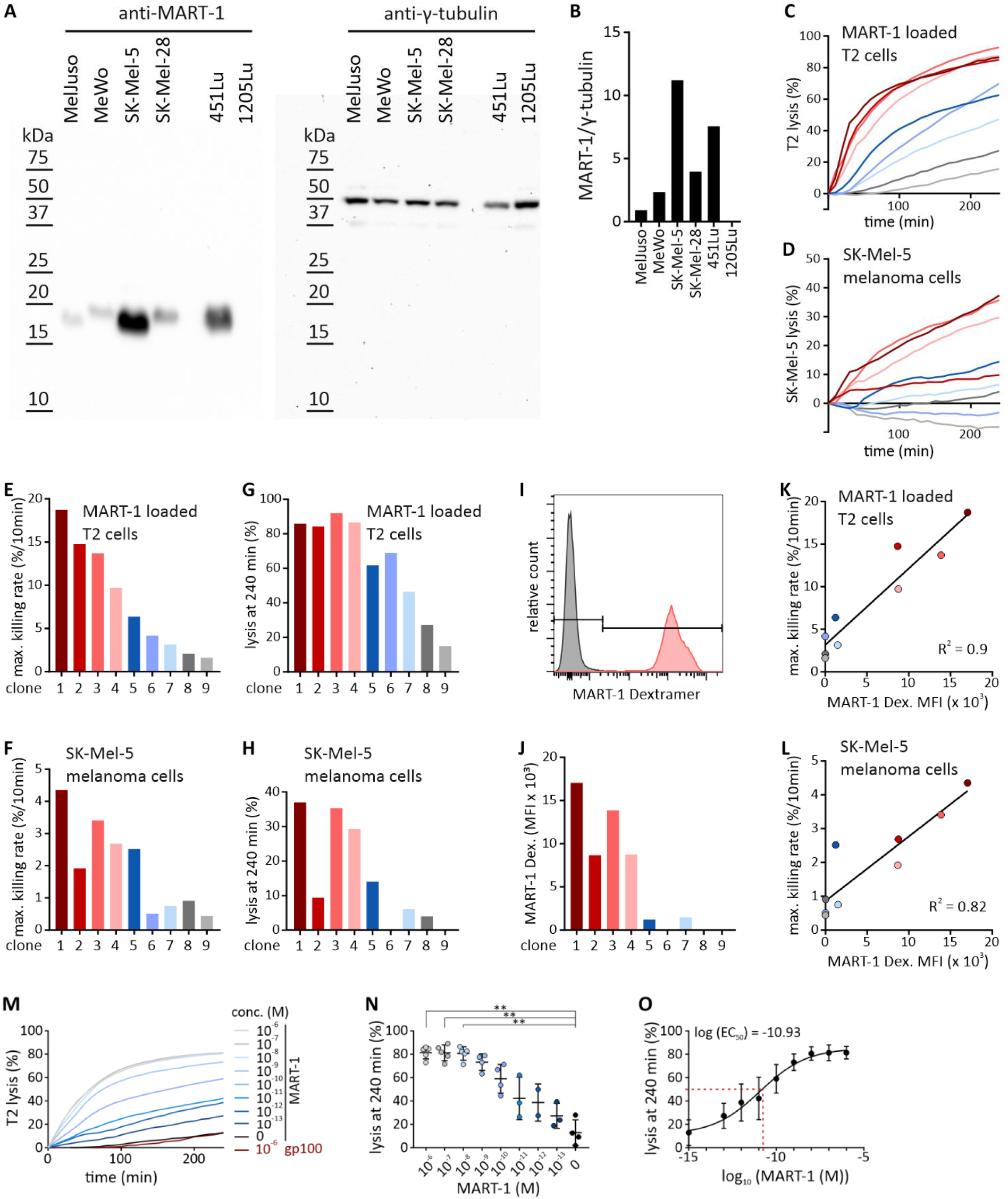
(A, B) MART-1 expression of different melanoma cell lines and functional analysis of MART-1-specific primary human CTL clones. Western blot analysis of different melanoma cell lines (A) and densitometric quantification against γ-tubulin (B). Marker ran in the empty lane between SK-Mel-28 and 451Lu. (C, D) MART-1-specific primary human CTL clones were generated using a modified protocol by Wölfl and Greenberg (Wolfl & Greenberg, 2014). Different CTL-MART-1 clones show different cytotoxicity against MART-1 peptide-loaded T2 cells (CTL:target ratio = 10:1, A) or against SK-Mel-5 melanoma cells (CTL:target ratio = 20:1, B), the latter of which present endogenous MART-1 antigen. (E, F) The maximal killing rate of the CTL-MART-1 clones against MART-1-loaded T2 cells and SK-Mel-5 cells was quantified. (G, H) Target cell lysis was quantified after 240 min for MART-1-loaded T2 cells and for SK-Mel-5 cells. (I, J) Antigen specificity was quantified using MART-1-specific dextramers in flow cytometry. One example (I) and quantification for all clones (J) are shown. (K, L) Maximal killing rates of CTL-MART-1 clones against MART-1-loaded T2 cells or SK-Mel-5 cells are correlated with the MART-1 dextramer mean fluorescence intensity (MFI). (M) To determine functional avidity of CTL-M3, real-time killing assays were performed with T2 cells loaded with different MART-1 antigen concentrations at an CTL-M3:SK-Mel-5 ratio of 5:1 (n=2-5). (N) Endpoint lysis at 240 min was quantified and plotted against the corresponding peptide concentration. (O) Data were fitted in a four parameter Hill equation revealing a log (EC_50_) of −10.93 M.

Fig. 1C-L illustrate cytotoxicity of 9 representative clones and their TCR-specificity against MART-1_26-35A27L_ using dextramer technology. These 9 clones from one of the 5 cloning approaches were chosen because their cytotoxicity against MART-1 loaded T2 cells (Fig. 1C) covered the full range from about 90% target elimination over 4 hours to less than 20%. We also tested the cytotoxicity of the 9 CTL clones against SK-Mel-5 melanoma cells. Compared to MART-1-loaded T2-cells, the 9 clones were less efficient against SK-Mel-5 cells (Fig. 1D) but the relative cytotoxicity of the 9 clones against their targets was similar for MART-1-loaded T2 cells and SK-Mel-5 cells. To quantify cytotoxic efficiency against MART-1-loaded T2 cells and SK-Mel-5 cells, we determined maximum killing rates (Fig. 1E, F) and lysis of targets cells at 240 min (Fig. 1G, H).

In addition, TCR-specificity of the 9 clones against MART-1_26-35A27L_ was quantified using dextramer technology. Fig. 1I shows an example of flow cytometry analysis, which was performed and quantified for all 9 clones (Fig. 1J). Among the clones, we found a correlation of MART-1-specific TCR expression and cytotoxic efficiency against MART-1-loaded T2 (Fig. 1K) or SK-Mel-5 (Fig. 1L) cells. In conclusion, we successfully generated different CTL-MART-1 clones, whose MART-1_26-35A27L_-specificities correlate well with their respective cytotoxic efficiency.

Because of its good cytotoxicity against MART-1-loaded T2 and SK-Mel-5 cells, CTLMART-1 clone 3 was analysed further and rigorously tested for stable cytotoxic efficiency during freezing/thawing cycles. Comparison of the cytotoxic efficiency for different clone expansions after several freezing/thawing cycles revealed a rather stable overall cytotoxic efficiency against MART-1-loaded T2 cells and even more stable against SK-Mel-5 cells. Therefore, clone 3 was used throughout the study and is named CTL MART-1-specific clone 3 (CTL-M3) from now on.

We next characterized the functional avidity of CTL-M3 against T2 cells loaded with different antigen concentrations. Very low cytotoxicity was observed if no antigen was present or if T2 cells were loaded with another common, albeit “wrong” melanoma-specific antigen, gp100 (Fig. 1M, two lowest lines). Cytotoxicity kinetics (Fig. 1M) and analysis of the endpoint lysis of target cells (Fig. 1N) demonstrated that - in a certain range - elimination of T2 cells by CTL-M3 was highly dependent on MART-1 antigen concentration. At 10^−8^ M and higher antigen concentrations, cytotoxicity and endpoint lysis did not change anymore, indicating saturation beyond 10^−8^ M. A fit of the endpoint lysis with a sigmoidal function revealed a half-maximal antigen concentration of about 10^−11^ M (Fig. 1O). In summary, we established robust MART-1 antigen-specific CTL which eliminate MART-1-loaded target cells in an antigen dose-dependent manner.

### At high effector-to-target ratios, CTL-M3 or NK cells completely eliminate SK-Mel-5 cells

A positive prognosis of many cancers correlates with the number of CTL which infiltrated the cancerogenic tissue (Fridman *et al.*, 2017) and a similar correlation is predicted for NK cells (Cursons *et al.*, 2019). We thus tested if CTL-M3 or primary NK cells are in principle able to completely eradicate all SK-Mel-5 cells at a high effector-to-target ratio up to 10:1 or 20:1. We first analysed the survival of SK-Mel-5 cells following 24 hour co-culture with different CTL-M3 or NK ratios using a flow cytometry assay. Quantification of these data revealed that few if any SK-Mel-5 cells survived the CTL-M3 or NK cell co-culture (Fig. 2A). We furthermore inspected the cells in parallel by microscopy and did not detect any viable-looking SK-Mel-5 cells after co-culture with CTL-M3 or NK cells (Fig. 2B). To quantify this, we used a single cell apoptosis-necrosis assay previously established in our group (Backes *et al.*, 2018). SK-Mel-5 cells were transfected with a pCasper-GR construct (a FRET GFP-RFP construct with a caspase3-cleavable site). Viable cells appear orange and switch to green if apoptosis is induced, and lose fluorescence if necrosis is induced (Fig. 2C, D). An overview over the whole field of the SK-Mel-5-CTL-M3 co-culture is shown in Fig. 2C. An example of a successful kill of a SK-Mel-5 cell by a CTL-M3 cell (indicated by the white arrows, the CTL-M3 is difficult to see because a 4x objective is used to image the whole well simultaneously) is depicted in Fig. 2D. A stacked plot showing the proportion of viable, apoptotic and necrotic cells over time (called death plot) of all SK-Mel-5 cells from this well reveals that close to 100% of all viable SK-Mel-5 cells are eliminated after 18 hours at a 4:1 effector-to-target-ratio (Fig. 2E). That means that all initially viable cells (orange) are either apoptotic (green) or necrotic and lost their fluorescence (grey). We could not recover any viable cells after 3-4 days co-culture, indicating that a high number of CTL-M3 are able to eradicate all SK-Mel-5 cells. We repeated the same approach with NK cells at an effector-to-target ratio of 25:1. Again, one complete well was imaged (Fig. 2F) and a successful kill is presented, in this case by necrosis, indicated by the loss of fluorescence (Fig. 2G). At this high ratio, all SK-Mel-5 cells were either killed by necrosis or apoptosis after 2 hours (Fig. 2H). We noticed that both CTL-M3 and NK cells sometimes kill by apoptosis followed by secondary necrosis, as indicated by the smaller number of apoptotic (green) SK-Mel-5 cells towards the end of the experiment compared to an earlier time point between 1-2 hours (Fig. 2E, H). We conclude that CTL-M3 or NK cells both eradicate all SK-Mel-5 cells in the culture at high effector-to-target ratios. This finding implies that there are no resistant SK-Mel-5 cells present in the initial culture.

**Fig. 2:**
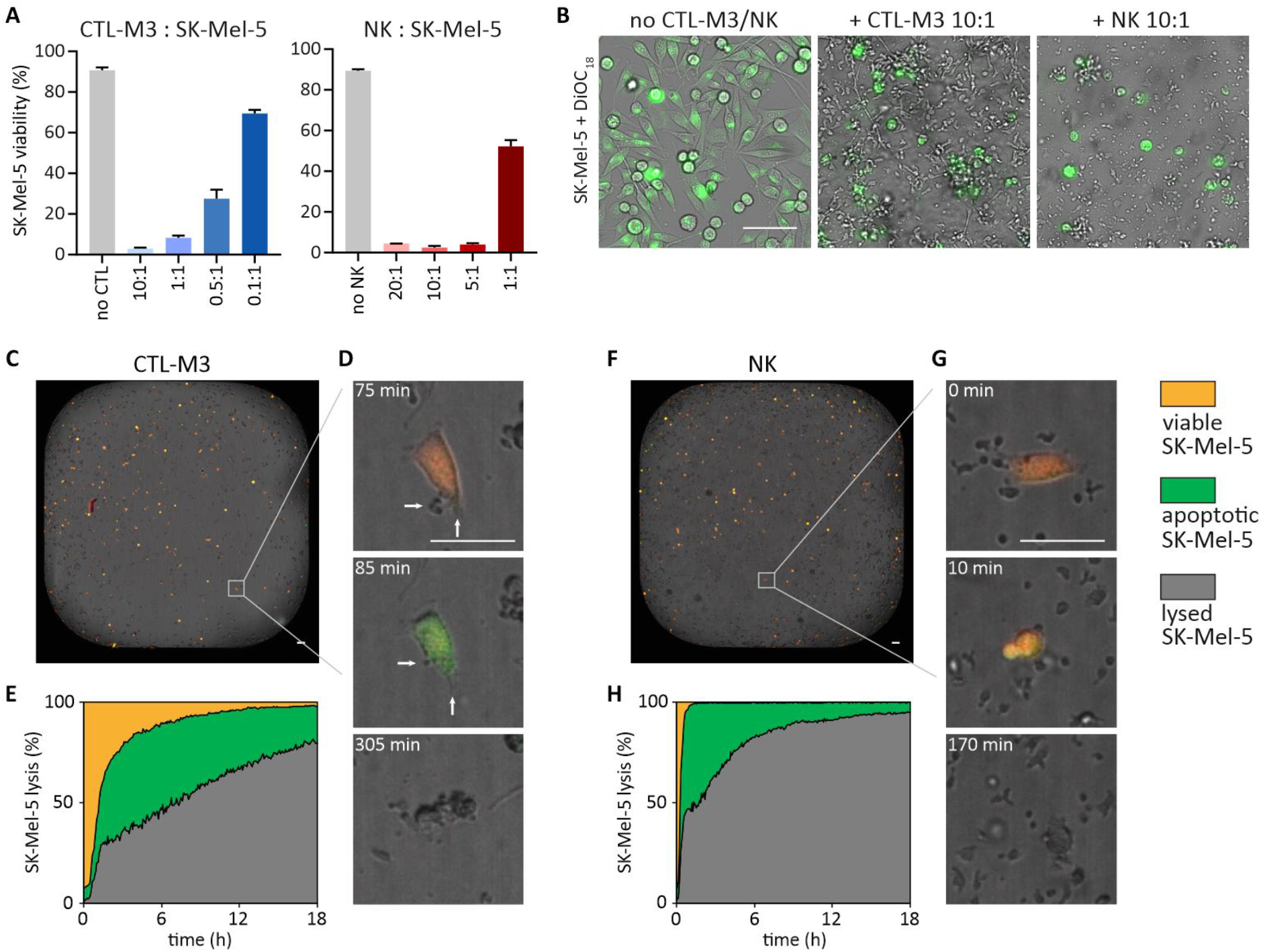
CTL-M3 or NK cells both completely eradicate SK-Mel-5 cells at high effector-to-target ratios. (A) Flow cytometry analysis of CTL-M3 or NK cell cytotoxicity at different effector-to-target ratios, respectively, against SK-Mel-5 cells. (B) Images of 5 × 10^4^ DiOC_18_-labelled SK-Mel-5 cells co-cultured with CTL-M3 (effector-to-target ratio was 10:1) or NK cells (effector-to-target ratio 10:1) for 24 h in a 48-well plate. (C-H) Real-time killing assays with 2 × 10^3^ transiently Casper-GR-transfected SK-Mel-5 cells in a 384-well plate. The whole field of view with all cells was analysed with the high-content imaging system. Overview of the complete well of SK-Mel-5 cells transiently transfected with Casper-GR incubated with CTL-M3 cell at an effector-to-target ratio of 4:1 (C-E) or with NK cells at an effector-to-target ratio of 25:1 (F-H). Example of successful killing of a SK-Mel-5 cell by a CTL-M3 (D, white arrows) or NK cell (G). Death plot of SK-Mel-5 cells of the whole well in the presence of CTL-M3 (E) or NK cells (H). Scale bars: 100 μm.

### A co-culture assay to analyse combined CTL-MART-1 and NK cell cytotoxicity against melanoma

Unfortunately, it seems unlikely that effector-to-target ratios of CTL or NK to cancer cells of 10:1 or higher are present in cancer tissue in the human body. Instead, lower effector-to-target ratios have been reported for NK cells in melanoma lesions (Balsamo *et al.*, 2012). Moreover, CTL and NK cells do not fight cancer independently of each other in the human body. To analyse combined CTL-M3 and NK cell cytotoxicity against the same cancer, i.e. melanoma SK-Mel-5 cells, we designed a suitable experimental setup (Fig. 3A, B). We established a co-culture assay, in which irradiated NK or CTL-M3 cells were used to avoid their further proliferation during co-culture. Irradiated CTL-M3 or NK cells kill as efficiently as their non-irradiated counterparts (Fig. 3C, D). SK-Mel-5 cells were pre-exposed to irradiated CTL-M3 or NK cells during co-culture at different effector-to-target ratios for 3-4 days, respectively (Fig. 3A). Following this CTL-M3 or NK cell pre-exposure during co-culture, we quantified cytotoxicity of fresh CTL-M3 or NK cells against surviving SK-Mel-5 cells, resulting in four different possible experimental combinations (Fig. 3B): 1. NK cell cytotoxicity after NK cell pre-exposure during co-culture, 2. CTL-M3 cytotoxicity after NK cell pre-exposure during co-culture, 3. NK cell cytotoxicity after CTL-M3 pre-exposure during co-culture, 4. CTL-M3 cytotoxicity after CTL-M3 pre-exposure during co-culture. Cytotoxicity of fresh CTL-M3 or NK cells was finally quantified using a kinetic "real-time" killing assay (Kummerow *et al.*, 2014).

**Fig. 3:**
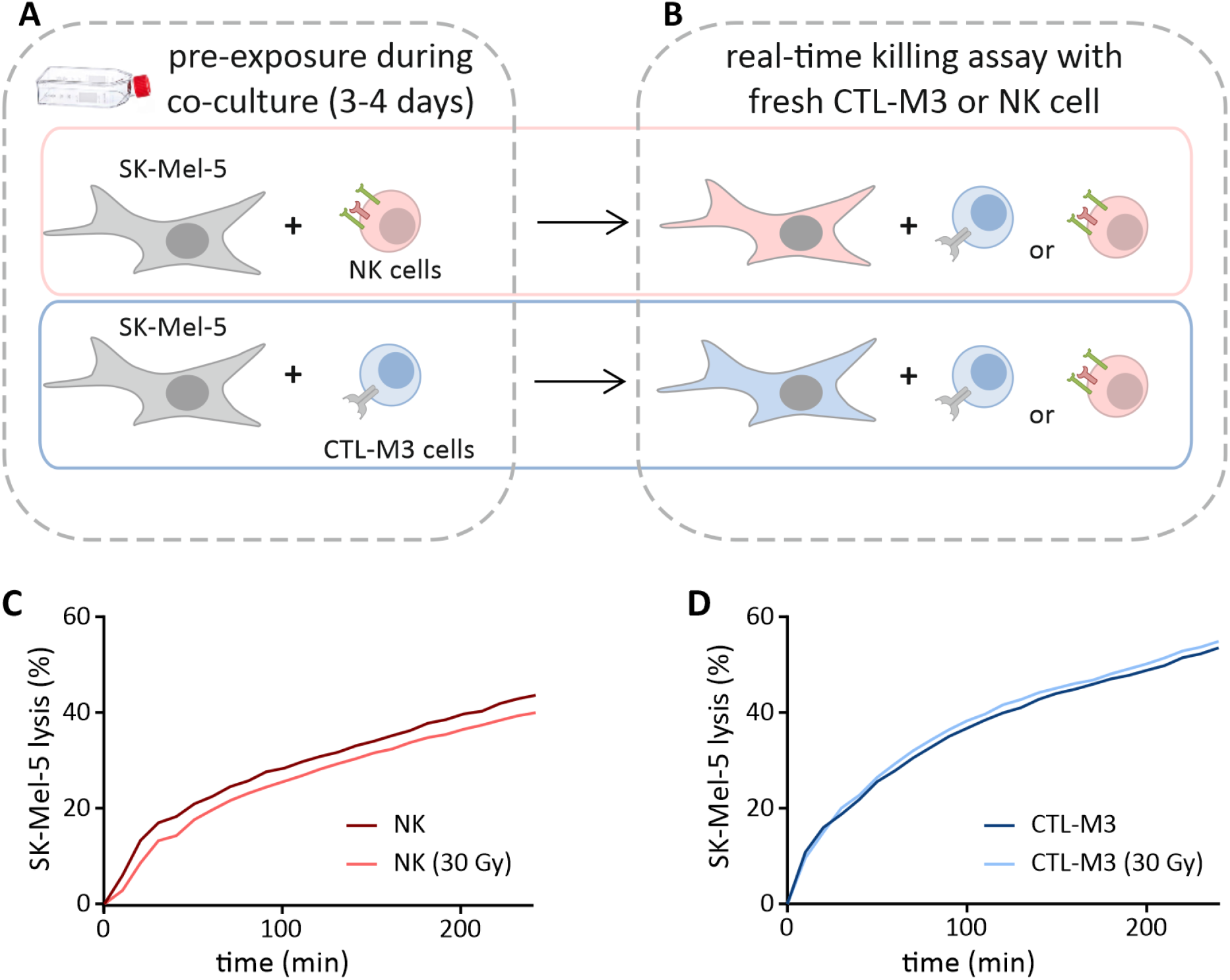
A co-culture assay to analyse the interdependence of CTL-M3 and NK cell cytotoxicity against SK-Mel-5 melanoma cells. (A) 10^6^ SK-Mel-5 cells were co-cultured for 3-4 days together with irradiated CTL-M3 or primary NK cells at different effector-to-target ratios, respectively. (B) SK-Mel-5 that survived CTL-M3 or NK cell pre-exposure, were harvested and then subjected to fresh CTL-M3 or NK cells and analysed by the real time-killing assay. Color-coded scheme: Orange-labelled cells depict SK-Mel-5 cells previously pre-exposed to NK cells; blue-labelled SK-Mel-5 cells depict SK-Mel-5 cells previously pre-exposed to CTL-M3 (C, D) Irradiated (30 Gy) CTL-M3 or NK cells eliminate SK-Mel-5 cells as efficiently as non-irradiated CTL-M3 or NK cells at an effector-to-target ratio of 10:1.

### NK and CTL-M3 cytotoxic efficiency against melanoma surviving NK cell pre-exposure

Applying the assay described in Fig. 3, we incubated NK and melanoma at effector-to-target ratios which did not eliminate all melanoma cells within 3-4 days. Subsequently, cytotoxicity of fresh CTL-M3 or NK cells against surviving SK-Mel-5 was quantified (Fig. 4, 5).

It was shown that melanoma cells acquired a protected phenotype against fresh human NK cells after surviving a long-term cocultures with low NK cell numbers (Balsamo *et al.*, 2012; Huergo-Zapico *et al.*, 2018). Testing NK cell cytotoxicity of fresh NK cells against melanoma cells surviving pre-exposure to insufficient NK cell numbers was done as proof of principle experiment. During the 3-4 day long primary encounter, NK:SK-Mel-5 cell ratios were varied between 2:1 and 8:1 including a control with no NK cells but NK-cell medium (Fig. 4A, B). Following this primary encounter, the surviving SK-Mel-5 cells were exposed to fresh NK cells at an NK:SK-Mel-5 ratio of 5:1, and cytotoxicity was analysed (Fig. 4B). Quantification of the maximal killing rate and the lysis at the end of the experiment revealed that the efficiency of fresh NK cells to eliminate surviving SK-Mel-5 was reduced at higher NK:SK-Mel-5 ratios during pre-exposure (Fig. 4B-D). Similar results were obtained in independent experiments if surviving SK-Mel-5 cells were exposed to fresh NK cells at a ratio of 10:1 (Fig. 4G-J). Despite clear trends of reduced NK cell cytotoxicity against pre-exposed SK-Mel-5 cells, the analysis did not reach statistical significance in most cases. We therefore re-tested the effect with another experimental approach with SK-Mel-5 cells from a different supplier at NK:SK-Mel-5 ratios between 2:1 and 10:1 during NK cell pre-exposure (Fig. 4K-N). While these SK-Mel-5 cells were killed by NK cells slightly less efficient (Fig. 4K), reduction of cytotoxic efficiency of fresh NK cells after pre-exposure was very similar (Fig. 4K-N), and statistical significance was reached in many cases. Cytotoxicity impairment of fresh NK cells against surviving SK-Mel-5 cells following NK cell pre-exposure is identical for both data sets (Fig. 4E). Considering the clear trends and statistical significances in many cases, we conclude that pre-exposure by low NK cell numbers induces resistance of SK-Mel-5 cells against a subsequent second NK cell exposure. It is very unlikely that resistant SK-Mel-5 cell pre-exist in the original culture, since all SK-Mel-5 cells were killed by sufficiently high numbers of CTL-M3 or NK cells (Fig. 2). We conclude that pre-exposure of NK cells results in the survival of SK-Mel-5 cells which likely acquire resistance towards cytotoxicity of freshly applied NK cells similar as found by Balsamo et al. (Balsamo *et al.*, 2012).

**Fig. 4:**
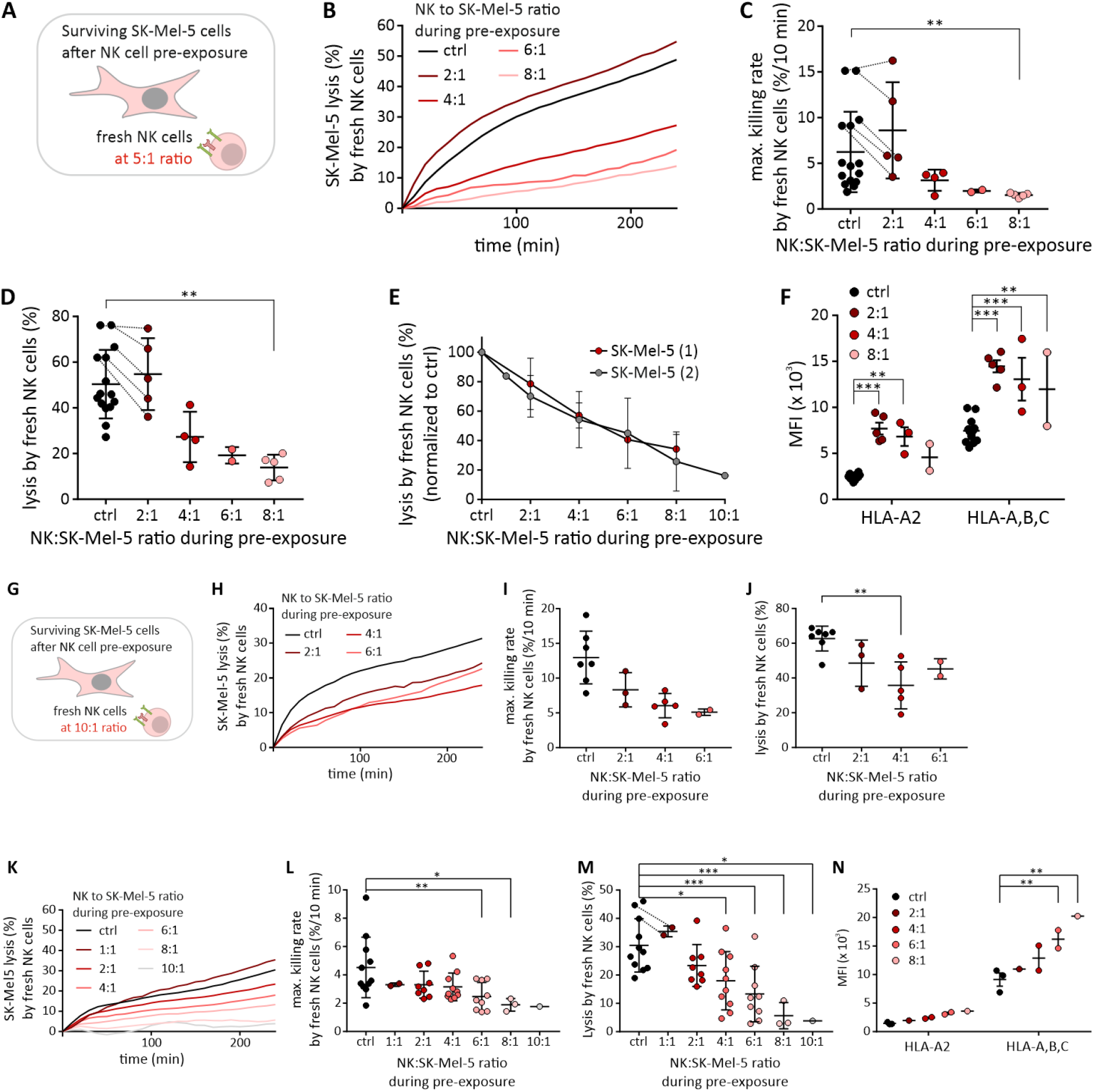
NK‒SK-Mel-5 pre-exposure induces resistance of surviving SK-Mel-5 cells against NK-mediated cytotoxicity. SK-Mel-5 cells were co-cultured for 3-4 days together with primary NK cells (A) at different NK:SK-Mel-5 ratios as indicated in (B). (B) NK cell-mediated cytotoxicity was tested against surviving SK-Mel-5 cells at a fixed NK:SK-Mel-5 ratio of 5:1 following different NK:SK-Mel-5 pre-exposure ratios (B). Maximal killing rate (C) and endpoint lysis after 240 min (D) were analysed to quantify NK cell cytotoxicity, n = 2-15. (E) Normalized NK cell cytotoxicity against SK-Mel-5 cells is compared between the data sets shown in Fig. 4D (SK-Mel-5 (1) from ATCC) and Fig. 4M (SK-Mel-5 (2) from CSL). Relative target cell lysis (normalized to killing with no NK cells present during pre-exposure) is displayed against the respective NK:SK-Mel-5 pre-exposure ratio. (F) HLA-A2 and HLA-A, -B, -C expression of SK-Mel-5 cells pre-exposed at different NK:SK-Mel-5 ratios were analysed by flow cytometry, n = 2-12. (G-J) Independent confirmation that NK‒SK-Mel-5 pre-exposure induces resistance against NK-mediated cytotoxicity. Same as (A), only that fresh NK cells were used at a ratio of 10:1. SK-Mel-5 cells were pre-exposed for 3-4 days to NK cells at different NK:SK-Mel-5 ratios as indicated in (H). (H) NK cell-mediated cytotoxicity which was tested against surviving SK-Mel-5 cells at an NK:SK-Mel-5 ratio of 10:1. Maximal killing rate (I) and endpoint lysis after 240 min (J) were analyzed to quantify NK cell cytotoxicity, n=2-7. (K-N) Second independent confirmation that NK‒SK-Mel-5 pre-exposure induces resistance against NK-mediated cytotoxicity. SK-Mel-5 (SK-Mel-5 (2)) cells from another supplier (CLS) were pre-exposed for 3-4 days to NK cells at different NK:SK-Mel-5 ratios (shown in K). (K) NK-mediated cytotoxicity against surviving SK-Mel-5 cells at an NK:SK-Mel-5 ratio of 5:1. Maximal killing rate (L) and endpoint lysis after 240 min (M) were analyzed to quantify NK-mediated cytotoxicity, n=1-11. (N), HLA-A2 and HLA-A, B, C expression of SK-Mel-5 cells pre-exposed to different numbers of NK cells were analyzed with flow cytometry, n=1-3.

**Fig. 5:**
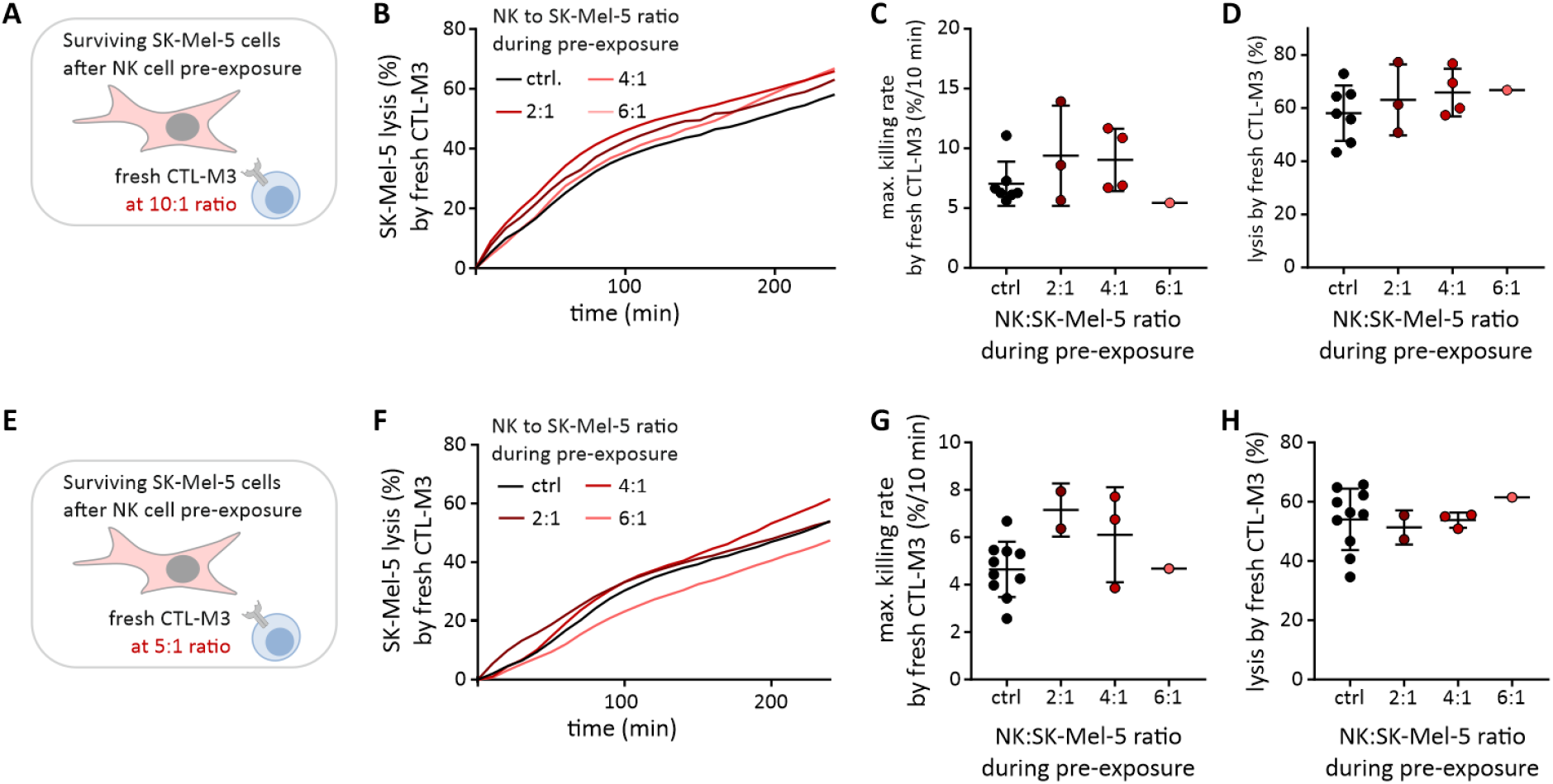
NK‒SK-Mel-5 pre-exposure does not induce resistance of surviving SK-Mel-5 cells against CTL-M3-mediated cytotoxicity. SK-Mel-5 cells were co-cultured for 3-4 days together with primary NK cells (A) at different NK:SK-Mel-5 ratios (B). (B) CTL-M3-mediated cytotoxicity against surviving SK-Mel-5 cells at a fixed 10:1 ratio following pre-exposure to NK cells at different ratios during co-culture. Maximal killing rate (C) and endpoint lysis after 240 min (D) were analysed to quantify CTL-M3-mediated cytotoxicity, n = 1-7. (E-H) Independent confirmation that NK‒SK-Mel-5 pre-exposure does not induce resistance against CTL-M3-mediated cytotoxicity. Same as (A), only that fresh CTL-M3 were used at a ratio of 5:1. SK-Mel-5 cells were pre-exposed for 3-4 days to NK cells at different NK:SK-Mel-5 ratios (F). (F) CTL-M3-mediated cytotoxicity against surviving SK-Mel-5 cells at a 5:1 ratio. Maximal killing rate (G) and endpoint lysis after 240 min (H) were analyzed to quantify CTL-M3-mediated cytotoxicity, n = 1-10.

One reason for this resistance of surviving SK-Mel-5 cells towards the cytotoxicity of fresh NK cells may be the upregulation of HLA class I molecules which are well-known as NK cell inhibitory receptors (Orr & Lanier, 2010; Balsamo *et al.*, 2012; Huergo-Zapico *et al.*, 2018). Their upregulation has been shown following INF-γ release by killer cells (Malmberg *et al.*, 2002; Derre *et al.*, 2006). We therefore tested HLA class I expression and indeed found upregulation of HLA-A2 and HLA-A, -B, -C in SK-Mel-5 cells incubated with NK cells (Fig. 4F, N). We conclude that NK–SK-Mel-5 co-culture reduces subsequent cytotoxic efficiency by fresh NK cells and that this effect could, in principle, be explained by HLA class I upregulation in SK-Mel-5 cells.

We next tested the cytotoxicity of CTL-M3 against surviving SK-Mel-5 cells (at a 10:1 CTL-M3:SK-Mel-5 ratio) after pre-exposure to NK cells (Fig. 5A). Fig. 5B shows that cytotoxicity of CTL-M3 was not reduced after NK cell pre-exposure at NK:SK-Mel-5 effector-to-target ratios between 2:1 and 6:1 during the co-culture compared to control. There was even a slight enhancement of CTL-M3 cytotoxic efficiency after NK–SK-Mel-5 co-culture (Fig. 5B) which is also evident from the analysis of the maximal killing rate (Fig. 5C, except the 6:1 pre-exposure condition) and the target cell lysis at 240 min (Fig. 5D). Independent experiments at an CTL-M3:SK-Mel-5 ratio of 5:1 confirmed that cytotoxicity after NK cell pre-exposure during the co-culture was not reduced under these conditions but even, at some ratios during pre-exposure, slightly enhanced at all NK-SK-Mel-5 co-culture conditions (Fig. 5E-H). In conclusion, cytotoxic efficiency after NK cell pre-exposure during co-culture is reduced for fresh NK cells but not (or on the contrary even slightly enhanced) for CTL-M3 cells against surviving SK-Mel-5 cells.

### NK and CTL-M3 cytotoxic efficiency against melanoma surviving CTL pre-exposure

To further examine the interdependence of NK cell and CTL-M3 cytotoxic efficiency, we reversed the approach and pre-exposed SK-Mel-5 melanoma cells with different CTL-M3:SK-Mel-5 ratios between 0.5:1 and 2:1 (Fig. 6A) and tested the cytotoxicity of fresh CTL-M3 or NK cells after CTL-M3 pre-exposure (Fig. 6, 7). The pre-exposure ratios between 0.5:1 and 2:1 were chosen because all SK-Mel-5 cells were eliminated at higher CTL-M3 to SK-Mel-5 ratios. NK cell cytotoxicity against surviving SK-Mel-5 cells (at 10:1 NK:SK-Mel-5 ratio, Fig. 6A) following CTL-M3 pre-exposure was already significantly reduced at very low CTL-M3:SK-Mel-5 ratios during pre-exposure (Fig. 6B) as quantified by the maximal killing rate (Fig. 6C) and the final lysis at 240 min (Fig. 6D). Independent experiments at 5:1 NK:SK-Mel-5 ratio confirmed that NK cytotoxicity after CTL-M3 pre-exposure was reduced (Fig. 6F-I). Pre-exposure of SK-Mel-5 cells to CTL-M3 clearly increased HLA class I expression (Fig. 6E), similarly as observed following NK–SK-Mel-5 pre-exposure (compare Fig. 4F, N). Increased HLA class I expression on SK-Mel-5 cells explains the reduced NK cell cytotoxicity after CTL-M3 pre-exposure.

**Fig. 6:**
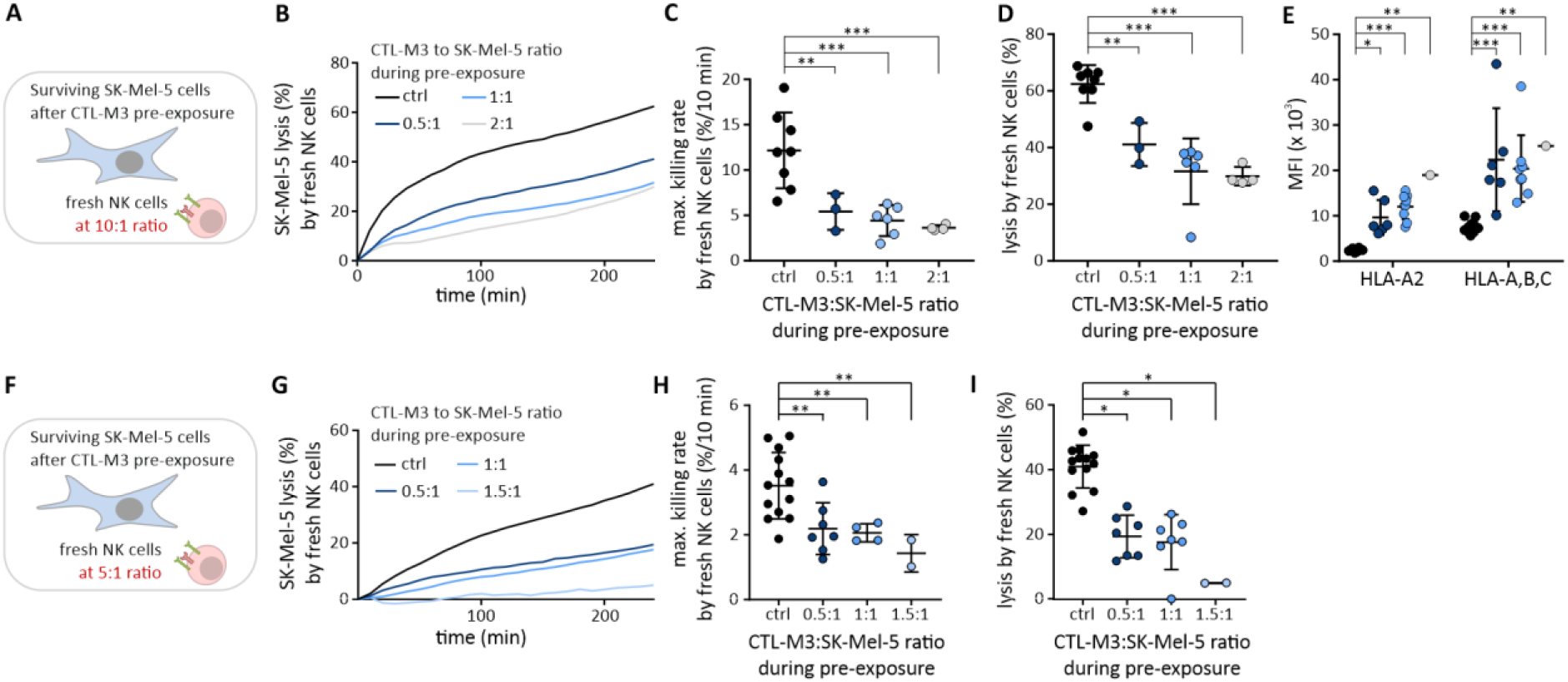
CTL-M3‒SK-Mel-5 pre-exposure induces resistance of surviving SK-Mel-5 cells against NK-mediated cytotoxicity. SK-Mel-5 cells were pre-exposed for 3-4 days to CTL-M3 (A) at different CTL-M3:SK-Mel-5 ratios (B). (B) NK cell-mediated cytotoxicity against surviving SK-Mel-5 pre-exposed to CTL-M3 cells, tested at a fixed NK:SK-Mel-5 ratio of 10:1 after different CTL-M3:SK-Mel-5 pre-exposure ratios. Maximal killing rate (C) and endpoint lysis after 240 min (D) were analysed to quantify NK cell-mediated cytotoxicity against SK-Mel-5 cells surviving pre-exposure, n = 3-8. (E) HLA-A2- and HLA-A, -B, -C expression of SK-Mel-5 cells pre-exposed to different numbers of CTL-M3 cells were analysed in flow cytometry, n = 1-12. (F-I) Independent confirmation that CTL-M3‒SK-Mel-5 pre-exposure induces resistance against NK-mediated cytotoxicity. Same as (A), only that fresh NK cells were used at a ratio of 5:1. SK-Mel-5 cells were pre-exposed for 3-4 days to CTL-M3 at different CTL-M3:SK-Mel-5 ratios (G). (G) NK cell-mediated cytotoxicity against surviving SK-Mel-5 cells. Maximal killing rate (H) and endpoint lysis after 240 min (I) were analyzed to quantify NK-mediated cytotoxicity against surviving SK-Mel-5 cells, n = 2-13.

**Fig. 7:**
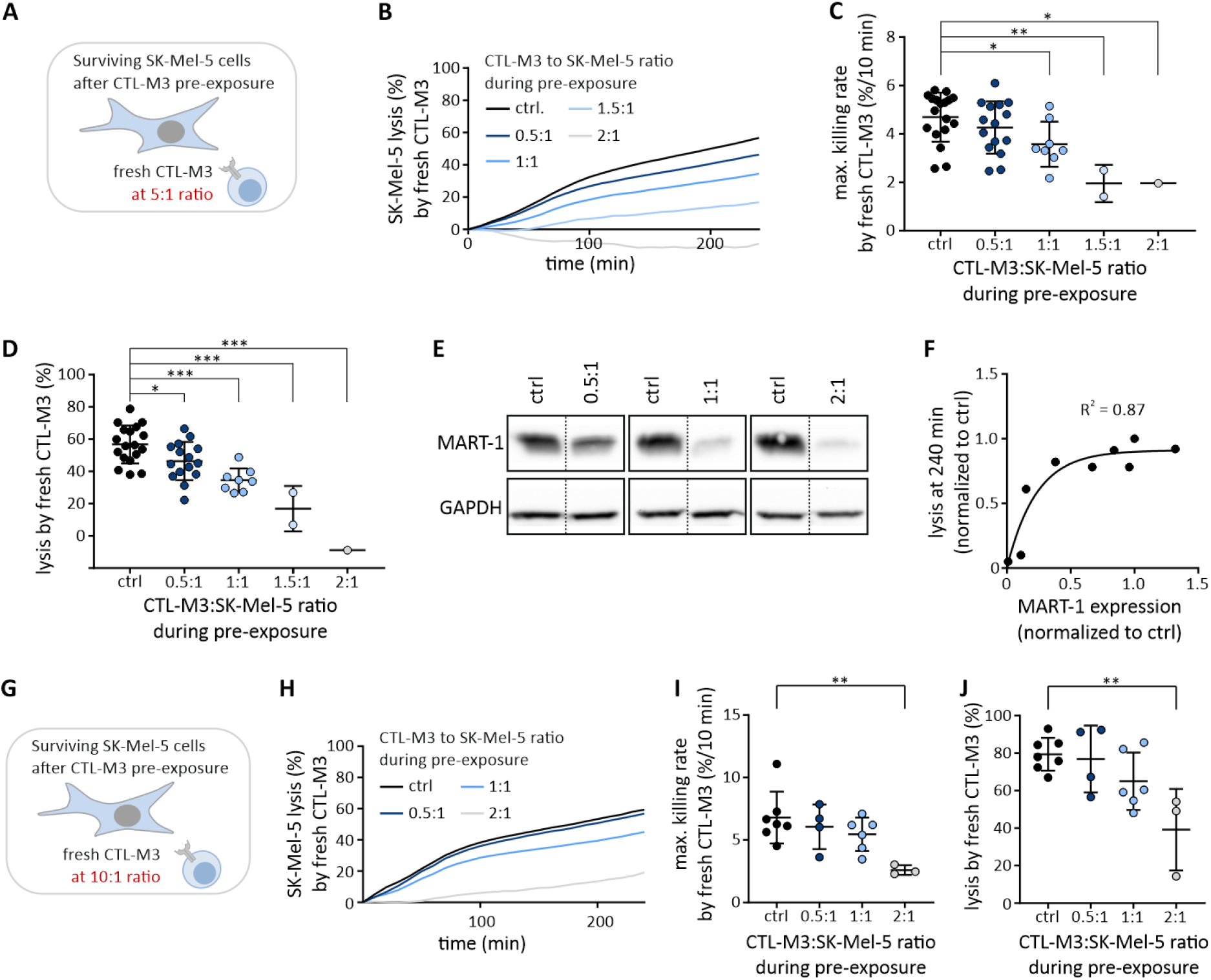
CTL-M3‒SK-Mel-5 pre-exposure induces resistance of surviving SK-Mel-5 cells against CTL-M3-mediated cytotoxicity. SK-Mel-5 cells were pre-exposed for 3-4 days to CTL-M3 (A) at different CTL-M3:SK-Mel-5 ratios (B). (B) CTL-M3-mediated cytotoxicity against surviving SK-Mel-5 cells was tested at a fixed CTL-M3:SK-Mel-5 ratio of 5:1 after different CTL-M3:SK-Mel-5 pre-exposure ratios. Maximal killing rate (C) and endpoint lysis after 240 min (D) were analysed to quantify CTL-M3-mediated cytotoxicity against surviving SK-Mel-5 cells, n = 1-18. (E) Western blots of SK-Mel-5 MART-1 expression after different CTL-M3‒SK-Mel-5 pre-exposure conditions next to control conditions (no CTL-M3 during pre-exposure). The full blots are shown in Fig. 8 with the respective conditions shown in Fig. 7E marked in red. The dotted line indicates that the bands were not juxtaposed on the blot, but they were always from the same blot. (F) Correlation of SK-Mel-5 lysis as measured in (B and D) against the respective MART-1 protein amount as measured in (E) or in Fig. 8. (G-J) Independent confirmation that CTL-M3‒SK-Mel-5 pre-exposure induces resistance against CTL-M3-mediated cytotoxicity. Same as (A), only that fresh CTL-M3 were used at a ratio of 10:1. SK-Mel-5 cells were pre-exposed for 3-4 days to CTL-M3 (G) at different CTL-M3:SK-Mel-5 ratios (H). (H) CTL-M3-mediated cytotoxicity against surviving SK-Mel-5 cells. Maximal killing rate (I) and endpoint lysis after 240 min (J) were analyzed to quantify CTL-M3-mediated cytotoxicity against surviving SK-Mel-5 cells, n = 3-7.

Considering increased HLA class I expression on surviving SK-Mel-5 cells following CTL-M3 pre-exposure, we predicted that fresh CTL-M3 should eliminate surviving SK-Mel-5 cells more efficiently after CTL–M3-SKMel5 pre-exposure (Fig. 7A). We tested this prediction by applying fresh CTL-M3 cells at an CTL-M3:SK-Mel-5 ratio of 5:1 subsequently to CTL-M3–SK-Mel-5 pre-exposure (Fig. 7A, B). However, to our surprise, cytotoxicity of fresh CTL-M3 was significantly reduced (Fig. 7B) as quantified by the maximal killing rate (Fig. 7C) and the endpoint target lysis at 240 min (Fig. 7D). Independent experiments at a 10:1 CTL-M3:SK-Mel-5 ratio confirmed that cytotoxicity after pre-exposure with CTL-M3 was reduced (Fig. 7G-J).

Since expression of HLA-A2 and HLA-A, -B, -C was increased on SK-Mel-5 cells (compare Fig. 6E) following CTL-M3 pre-exposure, reduced HLA class I expression was eliminated as a potential cause for reduced cytotoxicity of fresh CTL-M3 against surviving SK-Mel-5 cells. Another potential explanation for the reduced cytotoxic efficiency of fresh CTL-M3 against SK-Mel-5 cells pre-exposed to CTL-M3 could be a reduction of MART-1 antigen expression or presentation on surviving SK-Mel-5 cells. To test if reduced MART-1 expression of surviving SK-Mel-5 cells accounted for reduced CTL-M3 cytotoxic efficiency after CTL-M3-SK-Mel-5 pre-exposure, we tested MART-1 protein expression on SK-Mel-5 cells by western blot. Indeed, MART-1 expression was reduced compared to control after CTL-M3 pre-exposure on surviving SK-Mel-5 cells (Fig. 7E). Quantification of all conditions shown in Fig. 8 revealed a correlation between MART-1 expression of surviving SK-Mel-5 cells and CTL-M3 cytotoxicity against these SK-Mel-5 cells (Fig. 7F). Compared to the strong effect on MART-1 expression by CTL-M3 pre-exposure, NK cell pre-exposure only modestly reduced MART-1 expression at high effector-to-target ratios (Fig. 8). This latter finding is in line with the finding that CTL-M3 cytotoxicity against surviving SK-Mel-5 cells was not reduced but even slightly enhanced after NK cell pre-exposure (Fig. 5).

**Fig. 8:**
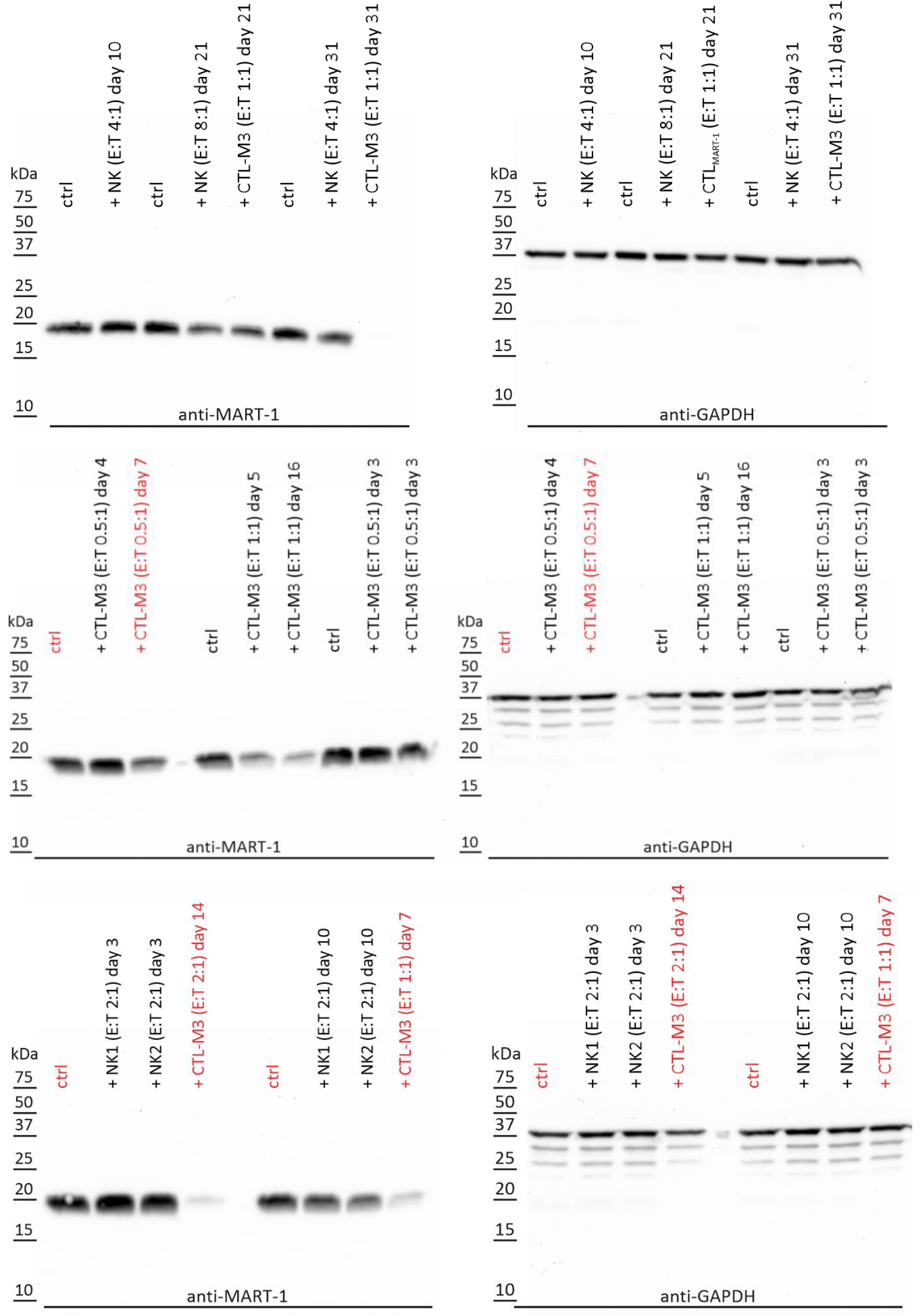
CTL‒M3-SK-Mel-5 pre-exposure induces down regulation of MART-1 protein expression on SK-Mel-5 cells. Complete Western Blots used in Figs. 7E and F. SK-Mel-5 cells were pre-exposed to CTL-M3 or NK cells at different E:T ratios. Indicated in red are the samples shown in Fig. 7E.

Considering these results, it is likely that reduction of MART-1 expression is responsible for reduced CTL-M3 cytotoxicity against surviving SK-Mel-5 cells after CTL-M3 pre-exposure. To test this hypothesis, we designed a rescue experiment by loading external MART-1_26-35A27L_ peptide on the surviving SK-Mel-5 cells. Fig. 9A illustrates the settings of the rescue experiment. Pre-exposure of CTL-M3 increases HLA-A2 and HLA-A, -B, -C expression on SK-Mel-5 cells (compare Fig. 6E) but decreases MART-1 expression (Fig. 7E, F). Thus, fresh CTL-M3 show reduced cytotoxicity against surviving SK-Mel-5 cells (Fig. 7B-D) which should be rescued by exogenous MART-1 antigen. We found that additional loading with MART-1 antigen on surviving SK-Mel-5 cells (Fig. 9A, right panel) was indeed sufficient to rescue CTL-M3 cytotoxicity against surviving SK-Mel-5 cells (Fig. 9B) as quantified by the maximal killing rates (Fig. 9C) and the endpoint lysis at 240 min (Fig. 9D).

**Fig. 9:**
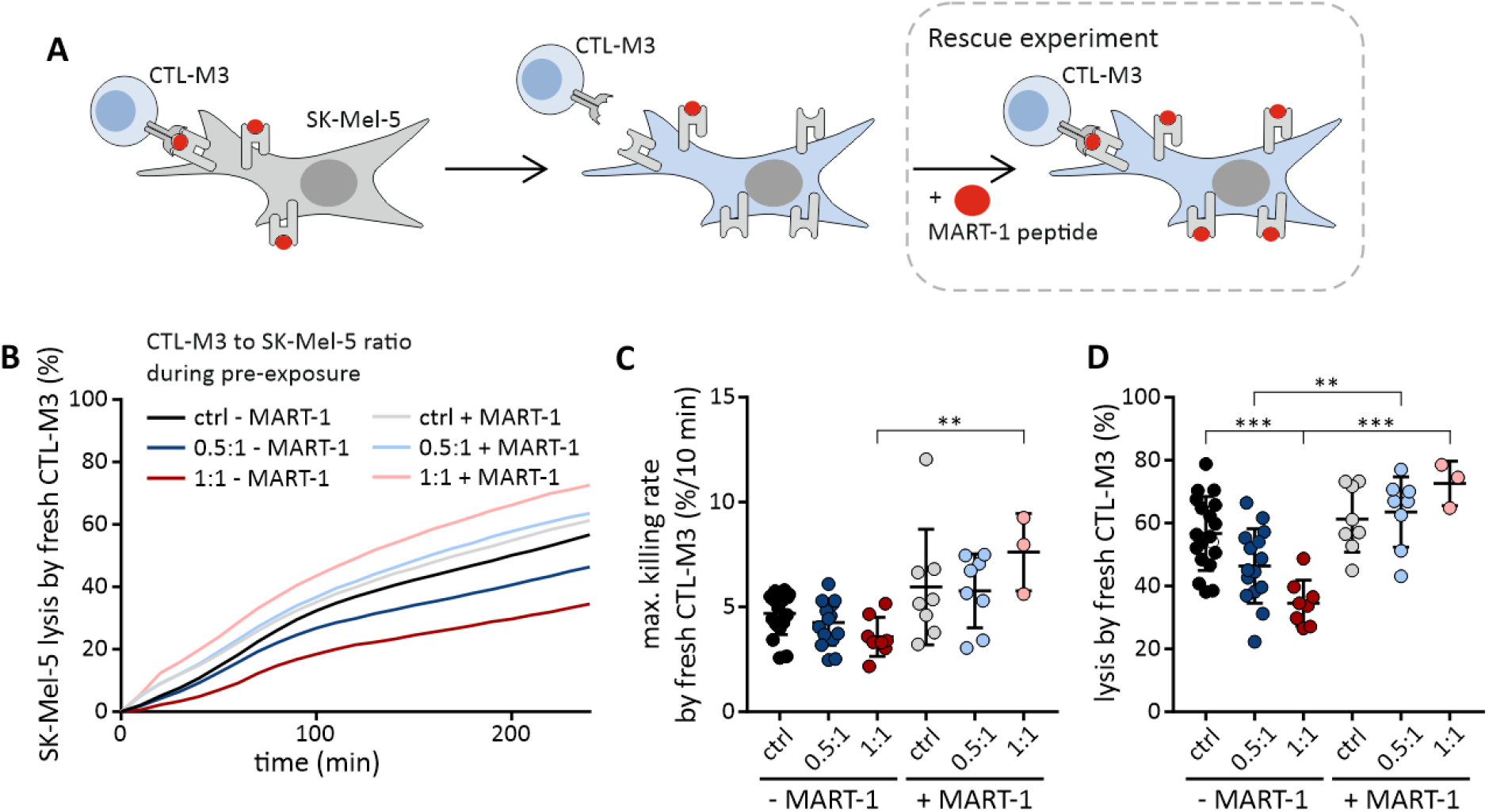
Susceptibility against CTL-M3 cytotoxicity can be rescued by loading surviving SK-Mel-5 cells with exogenous MART-1 peptide. (A) schematic representation of the experiment. SK-Mel-5 cells surviving CTL-M3 pre-exposure acquire resistance against CTL-M3 cytotoxicity by lowering MART-1 antigen expression while upregulating HLA class I molecules. Rescue strategy by exogenously loading of MART-1 antigen on SK-Mel-5 surviving CTL-M3 pre-exposure. (B) CTL-M3-mediated cytotoxicity against SK-Mel-5 cells at a fixed CTL-M3:SK-Mel-5 ratio of 5:1 measured after CTL-M3‒SK-Mel-5 pre-exposure, with (+ MART-1) or without (-MART-1) exogenous peptide loading. Control SK-Mel-5 (no co-culture with CTL-M3) were also included in the measurement. Maximal killing rate (C) and endpoint lysis after 240 min (D) were analysed to quantify CTL-M3-mediated cytotoxicity, n = 3-18.

The rescue experiments even induced more efficient CTL-M3 cytotoxicity compared to control conditions, which can be explained by the increased HLA-A2 expression on SK-Mel-5 cells following CTL-M3 pre-exposure and thus an increased MART-1-presentation on the surface of surviving SK-Mel-5 after peptide-loading.

We conclude that decreased MART-1 antigen expression on surviving SK-Mel-5 cells after pre-exposure to CTL-M3 cells is causative for reduced CTL-M3 cytotoxicity against the surviving SK-Mel-5 melanoma cells. Furthermore, enhanced HLA-A2 expression on surviving SK-Mel-5 cells after CTL-M3 pre-exposure increases CTL-M3 cytotoxicity against surviving SK-Mel-5 cells beyond control levels if additionally, exogenous MART-1 antigen is provided.

## Discussion

While it is common knowledge that cancer cells can evade from the immune system, only very few publications have directly addressed the quantification of CTL or NK cell cytotoxicity. Balsamo et al. showed that melanoma cells acquired a protective phenotype against human NK cells during long-term cocultures with low NK cell numbers as reported in tumours (Balsamo *et al.*, 2012). HLA class I expression was increased in melanoma cells in case insufficient NK cell numbers were present to eradicate all melanoma cells (Balsamo *et al.*, 2012; Huergo-Zapico *et al.*, 2018). Kohlhapp et al. and Xu et al. analysed the roles of murine CTL and NK cells with the result that both cytotoxic cell types are involved in the eradication of melanoma (Xu *et al.*, 2004; Kohlhapp *et al.*, 2015), and in addition Le et al. found that both CTL and NK cells are important to control solid tumours in a novel patient-derived xenograft model (Le *et al.*, 2020). Neubert et al. developed a well-controlled co-culture assay for human melanoma and CTL to analyse future T-cell based immunotherapies (Neubert *et al.*, 2016). None of these studies, however, was focused on if and how CTL and NK cells influence each other’s cytotoxicity. The reason for this may be the technical complexity. Thus, an assay is needed to quantify combined antigen-dependent CTL and NK cell cytotoxicity.

The assay we developed here allows exactly this: Analysing MART-1 specific human CTL and NK cell cytotoxicity using the same melanoma cell targets. As proof of principle for our assay, we first confirmed the finding that pre-exposure of melanoma to insufficient NK cell numbers renders surviving melanoma cells less susceptible to further NK cell cytotoxicity (*Balsamo et al., 2012*; Huergo-Zapico *et al.*, 2018). Making use of our assay, we then tested all combinations. We found that NK cell cytotoxicity after pre-exposure of melanoma to either CTL or NK was reduced as was CTL cytotoxicity after CTL pre-exposure. In all three conditions, pre-exposure induced resistance in surviving melanoma cells. Resistance was higher if high effector-to-target ratios of CTL or NK cells to melanoma cells were used during pre-exposure. This means that if not eradicated completely, a higher number of CTL or NK cells induces a higher degree of resistance in the surviving melanoma cells. Interestingly, a recent computational model predicts that it is beneficial for controlling tumor growth to apply lower CTL numbers several times rather than the equivalent cell number only once (*Khazen et al., 2019*). In support of a recurrent therapy with low CTL numbers, we found that CTL-M3 have a good cytotoxic potential at very low antigen concentration, which fits well with the important finding that T cells can be activated by very few antigen/MHC contacts (*Huang et al., 2013*). In principle, each antigen-specific CTL should be able to kill even if very little antigen is presented. To optimize the perfect dose of CTL for immunotherapy, kinetic information on individual CTL would be helpful, something that our microscopy assays can provide.

Is it possible to distinguish whether melanoma with pre-existing resistance were selected or if resistance was induced? Our finding that all melanoma cells can be eliminated by large numbers of CTL or NK cells strongly suggests that pre-exposure induces resistance of melanoma cells rather than selecting melanoma cell with pre-existing resistance. Melanoma, albeit not an epithelial cancer, can also undergo an epithelial-to-mesenchymal transition (EMT)-like transition towards mesenchymal traits. EMT has been implicated in carcinogenesis and metastasis of various cancers (Mittal, 2018) and can be induced in melanoma cell lines by NK cell editing (Huergo-Zapico *et al.*, 2018). It can be elicited within hours (Miettinen *et al.*, 1994) or days (Huergo-Zapico *et al.*, 2018), thus certainly in less than 3-4 days which was the pre-exposure time in our culture. Phenotype switching can however not only occur by EMT mechanisms and may involve common switch inducers (via EMT or other mechanisms) like transcription factors including SNAIls, ZEBs, TWIST, or c-Jun, but also the melanocytic lineage marker MITF (microphthalmia-associated transcription factor) or beta-catenin interaction factors LEF1 and TCF4 (Li *et al.*, 2015; Mittal, 2018). Signaling pathways responsible for EMT include Wnt, Notch, TGF-β, among others (Dongre & Weinberg, 2019). While phenotype switching can facilitate metastasis, migration, and invasion, it is not clear if it always induces melanoma resistance to CTL or NK cells. Only few of the numerous associated changes in gene expression have been related to melanoma resistance. Among others, melanoma cells have been shown to protect themselves by decreasing the anti-tumor activity of NK cells through inhibition of NK activating receptors like NKG2D (Pietra *et al.*, 2012) or by neutralizing CTL cytotoxicity through increased lysosome secretion (Khazen *et al.*, 2016). Both mechanisms, however, most likely do not explain our phenotypes as we expose the surviving melanoma to fresh CTL or NK cells.

Among the most relevant surface receptors modulating CTL and NK cell cytotoxicity against cancer, HLA class I receptors have a distinguished role because of their dual function. They are required for CTL cytotoxicity but inhibit NK cell cytotoxicity. An increase of HLA expression was reported following INF-γ release by killer cells (Malmberg *et al.*, 2002; Derre *et al.*, 2006). HLA class I receptor expression was indeed increased following melanoma-NK cell co-culturing, and this is considered a main reason to induce melanoma resistance against NK cell cytotoxicity (Balsamo *et al.*, 2012; Huergo-Zapico *et al.*, 2018).

In addition, also CTL pre-exposure enhanced HLA-A2 and HLA-A, -B, -C expression which can explain the reduced NK cell cytotoxicity against melanoma surviving either CTL or NK cell pre-exposure. HLA upregulation correlated with decreased NK cytotoxicity, pointing to a higher resistance of surviving melanoma cells exposed to high numbers of CTL or NK cells. This finding again favours the computational model prediction to apply lower CTL numbers several times rather than the equivalent number of cells at once (Khazen *et al.*, 2019). Increased HLA-A2 and HLA-A, -B, -C expression can also explain why CTL cytotoxicity against surviving melanoma was not reduced but even slightly enhanced after NK cell pre-exposure. It could be beneficial to combine NK and CTL immunotherapy but only if started with NK cells. While increased HLA-A2 and HLA-A, -B, -C offers a good explanation for these three cases, it does not explain the fourth combination. Higher HLA-A2 and HLA-A, -B, -C expression after CTL pre-exposure should not inhibit cytotoxicity of fresh CTL against the surviving melanoma cells but rather increase it. Reduced CTL cytotoxicity against surviving melanoma following CTL pre-exposure is, however, explained by a fundamental difference between CTL and NK cell pre-exposure. Both increased HLA-A2 and HLA-A, -B, -C expression but only CTL pre-exposure diminished expression of MART-1 antigen at the same time. The MART-1 antigen loss is the key factor, as adding MART-1 antigen to melanoma pre-exposed to CTL could completely rescue the subsequent cytotoxic efficiency by fresh CTL.

Quantification of CTL cytotoxicity has revealed differences between the *in vivo* and *in vitro* situation. Whereas efficient cytotoxicity has been reported in *in vitro* settings (Purbhoo *et al.*, 2004; Mempel *et al.*, 2006), cytotoxicity was found to be less efficient *in vivo* (*Boissonnas et al., 2007*; Breart *et al.*, 2008; Engelhardt *et al.*, 2012; Halle *et al.*, 2016). The reason for this is currently not clear. Additive cytotoxicity of CTL has recently been shown to significantly influence cytotoxic efficiency (Weigelin *et al.*, 2020). Our *in vitro* approach may add to the understanding of the differences of cytotoxic efficiency between *in vivo* and *in vitro* conditions. While we also find very high cytotoxicity as reported by others *in vitro*, we can also mimic drastically reduced cytotoxicity as reported in the *in vivo* situation. CTL cell pre-exposure drastically reduces CTL cytotoxic efficiency. CTL are usually present in persisting tumours and we predict that their presence should contribute to low *in vivo* CTL cytotoxicity. Thus, *in vitro* and *in vivo* discrepancies can be reconciled by our findings.

To quantify kinetics of cancer cell eradication by combination of CTL and NK cells could therefore prove a helpful *in vitro* tool to test cytotoxic efficiencies and quantify CTL and NK cell numbers for immune therapy. Considering our results, we propose the following:

1. If available and control of side effects allows this, patients should be treated with a large concentration of CTL or NK cells because only this guarantees the avoidance of escape mechanisms. This is a trivial result and we are of course not the first ones to propose this.
2. If not possible to eradicate the tumor at once with a very high dose of CTL or NK cells, it may be beneficial to repeatedly stimulate with lower doses to avoid strong melanoma resistance as has been suggested by Khazen et al. (Khazen *et al.*, 2019) based on a computer model. Our experimental set-up may well suited to analyse, what is the relevant time interval to still benefit from repeated treatment with low doses while avoiding induction of resistance to cytotoxicity.
3. Antigen loss may be a severe complication during extended or repeated CTL therapies, and this should be considered during immune therapy.
4. If possible, CTL immune treatment should be employed after NK cell treatment, which is also the physiological order of events.

## Additional information section

### Data Availability

All data n < 30 is made available according to the guidelines. Mean, SD and N are always provided.

### Competing interests

The authors declare that they have no competing interests.

### Author contributions

KSF, CK, ECS and MH are responsible for conception or design of the work with help from all authors regarding certain aspects. KSF performed and analysed most experiments. AK contributed and analysed experiments of Figure 3 and flow cytometry data. SC designed the assay for SK-Mel-5-NK cell co-culture and was supervised by CK and IB. CH and GS helped with generation of MART-1-specific CTL clones. All authors analysed and/or interpreted certain parts of the work. SI helped with interpretation of data and editing of the manuscript. KSF and AK co-wrote the methods section with corrections from ECS and MH. MH wrote the paper with help from KSF, ECS, AK, SI and all other authors. all authors edited and discussed the manuscript and the analysis of the data.

All authors approved the final version of the manuscript and all authors agree to be accountable for all aspects of the work in ensuring that questions related to the accuracy or integrity of any part of the work are appropriately investigated and resolved. All persons designated as authors qualify for authorship, and all those who qualify for authorship are listed.

### Funding

This project was funded by grants from the Deutsche Forschungsgemeinschaft (DFG), SFB 1027 (project A2 to MH, project C4 to IB) and SFB 894 (project A1 to MH) and a grant from the Bundesministerium für Bildung und Forschung (BMBF), 031L0133 (to MH).

## Acknowledgements

We very much appreciate the help of Prof. Hermann Eichler and the Institute of Clinical Hemostaseology and Transfusion Medicine at Saarland University Medical Center for obtaining human blood cells. We gratefully acknowledge Carmen Hässig for cell preparation. We also thank Elmar Krause and Jens Rettig for allowing us to use the flow cytometric sorting facility. We thank all lab members for insightful discussions.

## Notes

### Competing Interest Statement

The authors have declared no competing interest.

